# In-situ lipid profiling of insect pheromone glands by Direct Analysis in Real Time Mass Spectrometry

**DOI:** 10.1101/2022.05.19.492602

**Authors:** Nicolas Cetraro, Joanne Y. Yew

## Abstract

Lipid pheromones play a significant role in the behavior and ecology of many insects. The characterization of pheromone structures is a significant challenge due to their low abundance and ephemeral nature. Here we present a method for the analysis of lipid molecules from single pheromone glands of *Drosophila melanogaster* (fruit fly) using Direct Analysis in Real Time mass spectrometry (DART MS). Our results reveal that DART MS analysis of single tissues generates reproducible, species-specific lipid profiles comprised of fatty acids, wax esters, diacylglycerides and triacylglycerides. In addition, the ion source temperature and application of a solvent wash can cause significant qualitative and quantitative changes in the mass spectral profile. Lastly, we show that untargeted chemical fingerprinting of the gland can be used to accurately categorize species according to phylogenetic subgroup or genotype. Collectively, our findings indicate that DART MS is a rapid and powerful method for characterizing a broad range of lipids in tissues with minimal preparation. The application of direct tissue DART MS will expand the “secretome” of molecules produced by pheromone glands. In addition to its direct relevance to chemical ecology, the method could potentially be used in pharmaceutical studies for the screening and detection of tissue-specific drug metabolites.

## Introduction

Lipid pheromones play a significant role in the behavioral and reproductive decisions of many insects ^1^. Considering both the beneficial and destructive impact of insects to agriculture ^2^ and their importance as vectors of disease, the identification of pheromones and their function has been an area of extensive research. The characterization of pheromones can be technically challenging due to the limited amounts that are produced (usually at or below pmol levels of individual compounds) and ephemeral nature of the molecules. Analysis by gas chromatography MS (GCMS) is the most widely used method of analysis for lipid pheromone classes including saturated and unsaturated hydrocarbons, acetate esters, epoxides, terpenoids, fatty acids, and alcohols ^3^. However, under standard conditions, GCMS does not identify many of the heavier and more polar components without derivatization. In addition, sample preparation often requires dissection, extraction, and pooling samples from multiple individuals. For example, the analysis of drosophilid (fruit fly) pheromones typically requires 5 – 7 individuals yielding ca. 0.1 – 1 nmols of total lipids ^4-6^. Purification and structural elucidation of lipid pheromones has required several thousand flies ^7, 8^. Recently, laser desorption ionization (LDI) and matrix assisted laser desorption ionization (MALDI) MS have been used to characterize and image insect pheromone and lipid profiles directly from tissue ^7, 9-11^. The operation of LDI MS under intermediate pressure (∼2 mbar) allows in-situ detection of a broad range of lipid classes without derivatization or the application of a chemical matrix^7, 9, 12, 13^. Finer spatial resolution can be achieved with MALDI MSI approaches but necessitates a multi-step preparation process that entails embedding, sectioning, and matrix application ^14^.

Here, we describe the application of direct analysis in real time MS (DART MS), a form of plasma-based ambient ionization, for pheromone gland analysis. DART MS ionization uses a heated gas stream for analyte desorption through Penning ionization of atmospheric water and subsequent gas-phase ionization of analytes ^15, 16^. The open air ionization geometry allows individual tissue samples to be analyzed with minimal sample preparation, thus minimizing artifacts, sample loss, and degradation ^17^. The ion source produces a relatively large analysis area, ranging from 5 – 40 mm^2^, depending on the distance between the MS inlet and sample and the exposure time ^18^. Although the spatial resolution is not suitable for fine scale spatial imaging of single cells, the configuration allows for rapid chemical profiling of entire insect glands with minimal preparation. We show here the use of DART MS to characterize the ejaculatory bulb (EB), a pheromone gland found in male *Drosophila*. Reproducible, species-specific profiles of fatty acids (FAs), wax esters, diacylglycerides (DAGs), and triacylglycerides (TAGs) can be generated from single pieces of tissue without derivatization or purification. In addition, experimental parameters such as the ion source temperature and the application of a solvent wash quantitatively and qualitatively alter the profile. Lastly, we show different drosophilid subgroups or flies of different genotypes can be differentiated and classified on the basis of untargeted chemical profiling of EBs.

## Results and Discussion

### Single gland profiling of D. melanogaster

To determine the suitability of DART MS for direct tissue analysis, we optimized sample preparation and analytical parameters using the ejaculatory bulb (EB) of *D. melanogaster*. The EB, located in the terminal portion of the male reproductive system, is a male pheromone-producing gland that is conserved throughout the *Drosophila* family ^13, 19-21^ (**Fig. 1A**). The EB lipid contents have been well-characterized by GCMS and LDI MS and thus, the tissue serves as a useful analytical standard. Individual EBs were dissected and placed on stainless steel mesh (**Fig. 1A**). Five to seven biological replicates were analyzed using an automated linear rail system that allows for uniform sample positioning and analysis time per replicate. Each gland was exposed to the helium plasma for approx. 8 sec. Considering the diameter of the helium beam (5 – 40 mm^2^) relative to the size of the gland (ca. 200 - 300 µm in diameter), the entirety of the tissue is exposed during analysis. Direct analysis of freshly dissected EBs revealed 2 major peaks corresponding to two previously identified sex pheromones: 11*Z*-Octadecen-1-ol acetate (*cis*-vaccenyl acetate; *m/z* 311.30) ^21, 22^ and (3*R*,11*Z*,19*Z*)-3-acetoxy-11,19-octacosdien-1-ol (CH503; *m/z* 465.43) ^7, 23^ (**Fig. 1B; Supp. Fig. 1 & 2**). Two fragments with *m/z* reflecting the loss of an acetate group from each pheromone (*m/z* 251.27 and *m/z* 405.41, respectively) are also detected and are likely the result of thermal decomposition. DART MS analysis of synthetic cVA and CH503 also shows the same loss of acetate in addition to CH503 dehydration (**Supp. Fig. 2**). Other components of the gland that were identified on the basis of exact mass measurements include fatty acids (FAs), DAGs, and TAGs (**Supp. Table 1**). The DAGs are detected mainly as dehydrated ions ([M + H - H_2_O]^+^) The loss of water observed is consistent with the ionization behavior of a DAG standard analyzed under identical conditions (**Supp. Fig. 3**). Overall, the total ion chromatograms and mass spectral profiles were uniform between biological replicates in terms of total intensity and composition (**Fig. 1C**; **Supp. Fig. 1**). The abundance of various chemical classes changed qualitatively during the exposure period – molecules below *m/z* 500 were more abundant in the initial 3 sec whilst DAGs and TAGs were ionized mainly after 3-6 sec of exposure to the DART source (**Fig. 1D**). Some of the lipids, particularly DAGs and TAGs, may be derived from the fat body, a storage organ that contains a large amount of lipid reserves and surrounds the internal organs of insects ^24^. We have previously been able to detect pheromones secreted from the gland without dissection by analyzing whole flies with DART MS ^25^. However, compared to dissected tissue, reproducible MS profiles are difficult to generate from whole flies due to variation in ion source placement. In addition, other compounds from the external cuticle (e.g., hydrocarbons) are likely to also be ionized making it difficult to identify gland-specific compounds.

**Figure 1.**
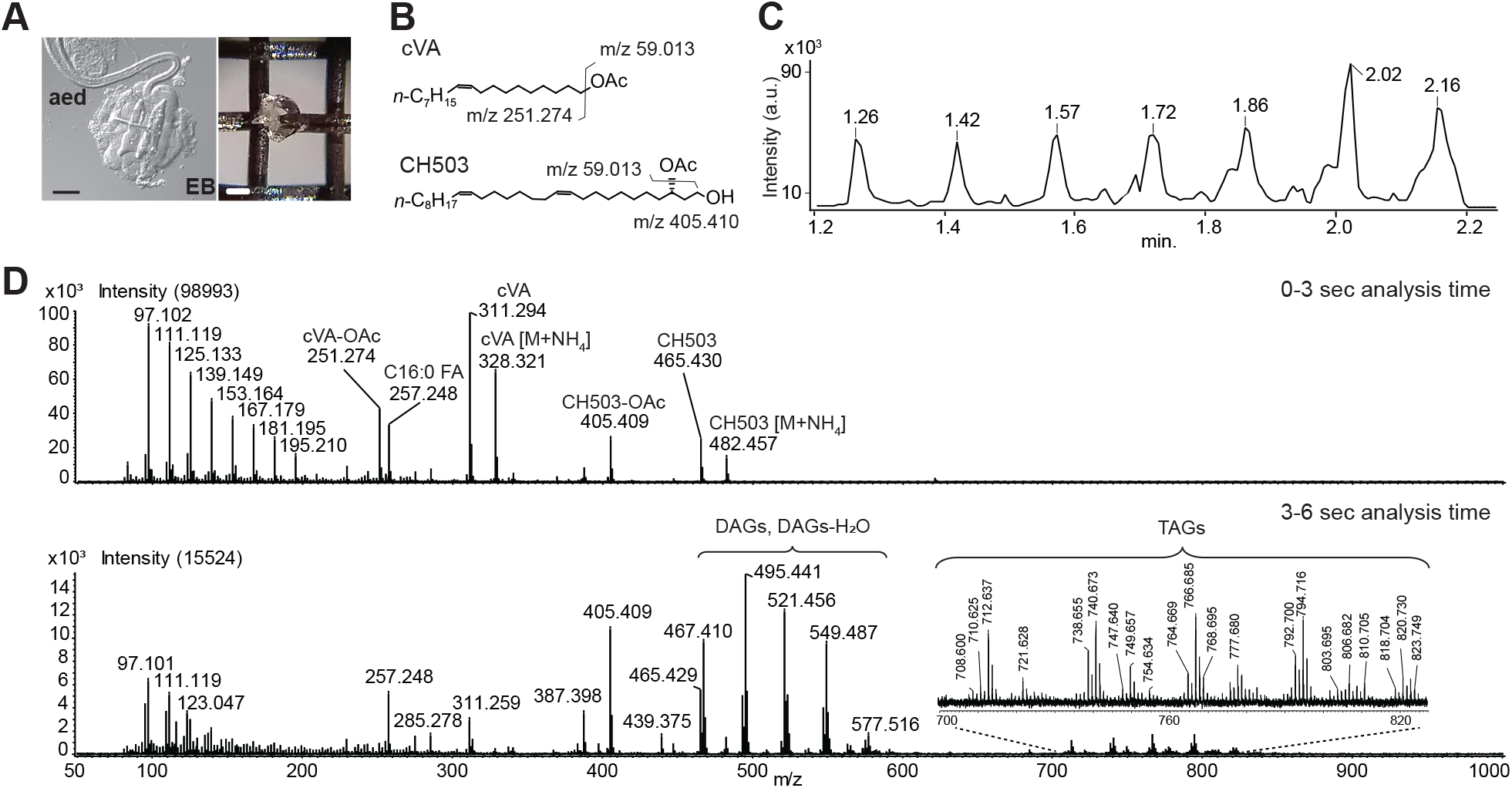
DART MS analysis of the ejaculatory bulb (EB). **(A)** Image of a single EB visualized under differential interference contrast illumination and placed on the stainless steel grid sample holder; aed: anterior ejaculatory duct; scale bar: 100 µm. **(B)** Chemical structures and predicted *m/z* of the two major pheromones produced in the EB: *cis*-vaccenyl acetate (cVA) and (3*R*,11*Z*,19*Z*)-3-acetoxy-11,19-octacosdien-1-ol (CH503). Fragments resulting from the loss of acetate groups are shown. **(C)** The total ion chromatogram from 7 EBs placed on a single QuickStrip shows relatively uniform intensity between each replicate; a.u.: arbitrary units. **(D)** Representative DART MS spectrum from a single gland in the first 3 sec of introduction to the ion source and the subsequent 3-6 sec of analysis. Mass signals corresponding to cVA, CH503, deacetylated fragments, fatty acids (FA), diacylglycerides (DAGs) and triacylglycerides (TAGs) are labeled. Profiles were acquired at 300 °C. Some of the FA and DAGs may be the products of TAG decomposition.

Thermal decomposition is a common occurrence in the DART source and has been observed for serum metabolites, sterols, and plant and animal oils ^26-28^; as such, some of the FA and DAG signals are also likely to be products of TAG decomposition. To differentiate endogenously produced molecules from ionization artifacts, we compared the EB profile generated with DART MS to *D. melanogaster* EB profiles previously obtained with laser desorption ionization MS ^20^. LDI MS is a soft ionization method with minimal fragmentation ^22^. As with DART MS analysis, the wax ester pheromones cVA and CH503 in addition to multiple species of FAs, DAGs, and TAGs were observed using LDI MS **(Supp. Table 1**). The detection of intact DAGs by LDI MS strongly supports the endogenous presence of these molecules in the EB but does not preclude the possibility that some of the DAG and FA signals observed in DART MS spectra may also be the products of thermal decomposition (see below).

### Effect of temperature on EB tissue profile

Increasing the temperature of the DART gas stream can significantly improve the ionization of heavier, more polar molecules but may also lead to the rapid degradation of smaller, more volatile compounds and increase the occurrence of ester hydrolysis ^26^. To characterize the impact of temperature on EB lipids, we analyzed dissected tissue over a range of DART source temperatures from 100 – 350 °C (**Fig. 2A**). Most compounds were reliably detected between 250 – 350 °C. Almost no signals were detected above baseline at temperatures below 150 °C. Increasing temperature resulted in a greater abundance of deacetylated cVA and CH503 (**Fig. 2B**). The slight increase in DAG levels at higher temperatures may be the result of increasing TAG decomposition. Our analysis of DAG and TAG standards confirmed that both lipid types lose acyl chains with increasing temperature (**Supp. Fig. 3**). We conclude that direct tissue analysis between 250 – 300 °C is optimal for detecting a breadth of lipid molecules. However, products arising from the decomposition of ester lipids such as tri-and diacylglycerides and acetate esters are inevitable and confound the differentiation of endogenous compounds from fragment artifacts without a secondary method. Nonetheless, with mass isolation capability, fragment ions can provide insight into the structural features and functional groups of lipids produced in biological tissues.

**Figure 2.**
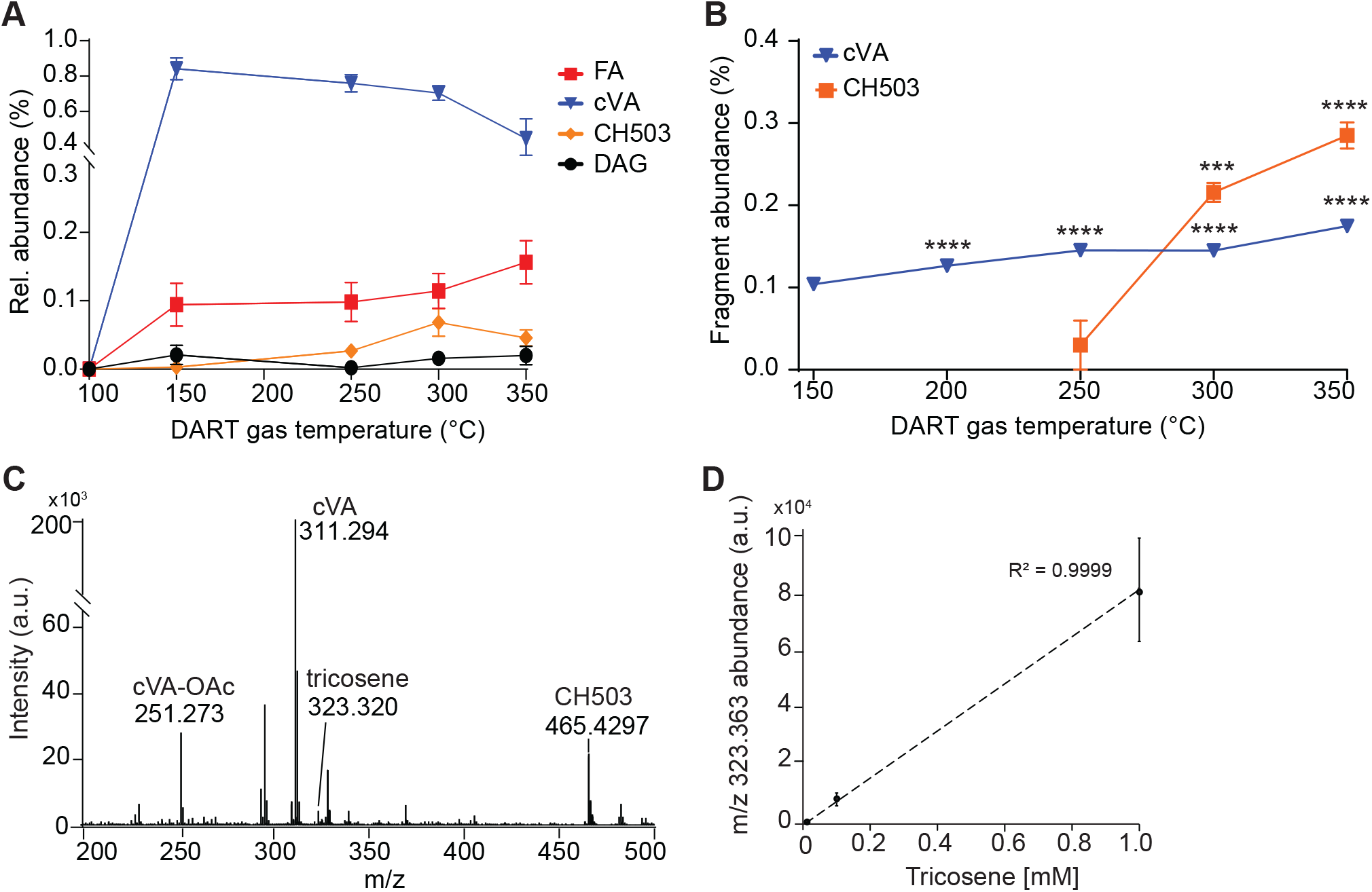
Effect of DART gas temperature on direct tissue analysis signal abundance and fragmentation. **(A)** Signals corresponding to wax ester pheromones, diacylglycerides (DAG) and fatty acids (FA) were detected reliably at 250 °C and above. **(B)** Fragmentation due to the loss of the acetate group from cVA or CH503 was significantly higher at 350 °C compared with 150 °C (cVA) or 250 °C (CH503); ***: p=0.0002, ****: p<0.0001, Holm-Sidak’s multiple comparisons test. **(C)** Representative DART MS spectrum measured at 300 °C from an EB spiked with 100 nmols tricosene. **(D)** The signal intensity of tricosene spiked onto the tissue increases linearly with concentration between 0.01 – 1 mM (R^2^ = 0.99). For all graphs, bars indicate mean ± S.E.M; n=5-7 EBs per point.

### Quantitative profiling of biological tissue

To investigate whether tissue profiling by DART MS can be used for quantitative analysis, we added an external alkene standard, tricosene (C23 H46; [M+H]^+^ 323.26), directly to dissected tissues at different concentrations (**Fig. 2C, D**). At an ion source temperature of 250 °C, the normalized signal intensity increased linearly with sample concentration between 0.01 mM to 1 mM (R^2^ = 0.99). Quantitative analysis is thus possible with the use of an appropriate internal standard with similar vapor pressure to the analytes of interest. However, considering the diverse array of endogenous lipid classes found within biological tissues, it will be necessary to use multiple standards over a range of vapor pressures in order to match the heterogeneity of compounds and their respective ionization efficiencies.

### Effect of solvent treatment on EB tissue profile

The use of a tissue wash in MALDI imaging protocols is an effective method for removing ion suppression by removing contaminants and excess salts ^29-32^. To test whether washing the tissue prior to DART MS analysis could improve ionization and overall signal intensity by removing salts and increasing membrane permeability, we applied solvents of various polarity to dissected glands and assessed the responses of four categories of molecules found in the gland. Our findings show that the area of the total ion chromatogram did not change significantly between solvent conditions, indicating that overall ion counts were similar between each condition (**Fig. 3A**). In contrast to no-solvent conditions, hexane and chloroform/ MeOH washes lowered the coefficient of variation between biological replicates. It may be the case that because biological tissue is heterogenous in its lipid composition and topology, the use of an apolar wash helps to homogenize lipid distribution, resulting in lower variation between tissue samples. Washing with hexane produced qualitative changes in the lipid profile - notably, DAG signal intensity increased significantly, though some of the signals may be the products of TAG fragmentation (**Fig. 3B; Supp. Fig. 4**). Overall, however, the use of solvents did not significantly improve sensitivity and in fact, worsened the outcome by increasing sample-to-sample variation or adding FA contamination. Analysis of solvent-only preparations showed FA signals, in some cases at levels indistinguishable from measurements with tissue samples (**Supp. Fig. 5**). Although the FA abundances of hexane-and MeOH-treated EBs were significantly higher than background levels, nonetheless, the contamination hinders interpretation and quantitation. In addition to FA contamination, EBs treated with EtOH exhibited a major signal at *m/z* 376.24 that was found in only at low levels in other conditions. Based on predicted elemental composition (C26 H34 O N) and high degree of conjugation (10 double bond equivalents), this molecule is likely to be a contaminant rather than an endogenous lipid. These results indicate that pre-treatment of the tissue with solvents provide little advantage for preliminary tissue profiling. Moreover, it will be critical to use ultrapure solvents and sample preparation tools when applying external standard solutions.

**Figure 3.**
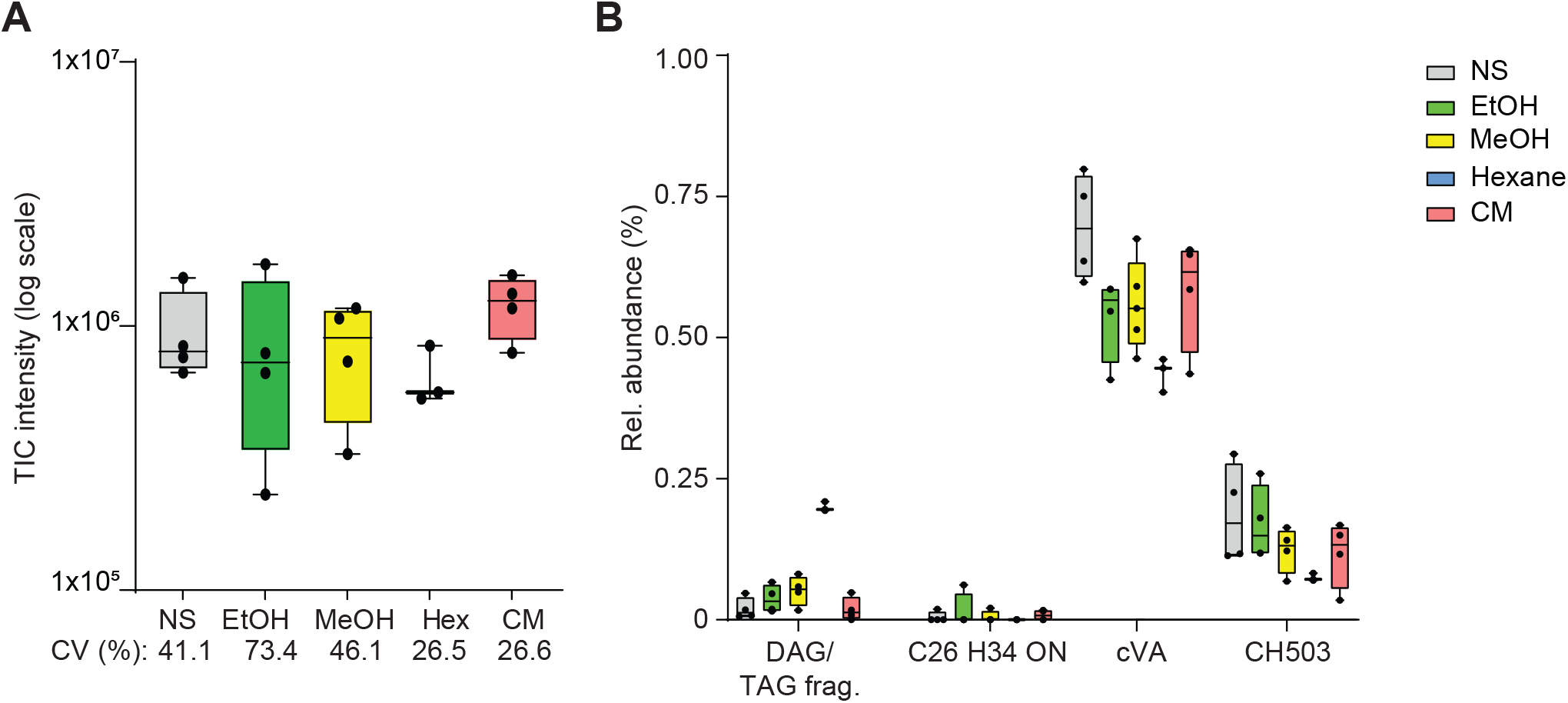
Addition of a solvent wash produces qualitative changes in DART MS profile. **(A)** The area beneath the total ion chromatogram did not change significantly between each of the solvent conditions (ANOVA, p=0.3773). The coefficient of variance (CV) is shown for each solvent condition; NS: no solvent, EtOH: ethanol, MeOH: methanol, Hex: hexane, CM: chloroform/ methanol. **(B)** The relative abundance of various components of the EB profile change with solvent conditions. The DAG abundance increased significantly with hexane application whilst levels of an unknown compound (C26 H34 ON), likely a contaminant, was most evident with EtOH wash. Hexane produced the least variability between replicates. The mean with maximum and minimum values is shown in each plot; n=3-5 per condition. All spectra were acquired at 300 °C.

### Classification by EB chemical profiles

Pheromone and lipid profiles can evolve rapidly between species ^33^. For this reason, chemotaxonomy using fatty acid profiles and other lipid features has been successfully employed for the classification of bacterial and animal species ^34-36^. Cuticular lipids have also been proposed as a useful trait for taxonomic classification, especially between closely related species with no obvious morphological differences ^37, 38^. We previously showed that the cleavage products of unsaturated fatty acids induced by ozonolysis combined with DART MS can be used for effective species classification ^35^. To test whether the EB chemical profiles generated by DART MS can be applied to taxonomic classification, we used principal components analysis (PCA) to classify EB profiles from 18 drosophilid species either belonging to the Hawaiian *Drosophila* picturewing clade, repleta group, or melanogaster group. The MS analyses from each species EB revealed a broad range of compounds ranging from *m/z* 150 – 705 (**Fig. 4A**). The members of the picturewing clade could be separated from other phylogenetic groups with 100% accuracy (verified with LOOCV) using PCA with spectral profiles. These findings indicate that endogenous lipid profiles serve as a useful phenotypic feature for classifying insects and inferring phylogenetic relationships. Many of the observed ions are novel molecules that have been missed by GCMS and are reported for the first time here. Chemical classification will require further structural elucidation by MS/MS. Moreover, in the absence of chromatographic separation and mass isolation, some single peaks are likely to represent isobaric molecules. Nonetheless, our results show that DART MS-generated chemical profiles can be used as taxonomic markers to distinguish invertebrate species even in the absence of complete structural elucidation.

**Figure 4.**
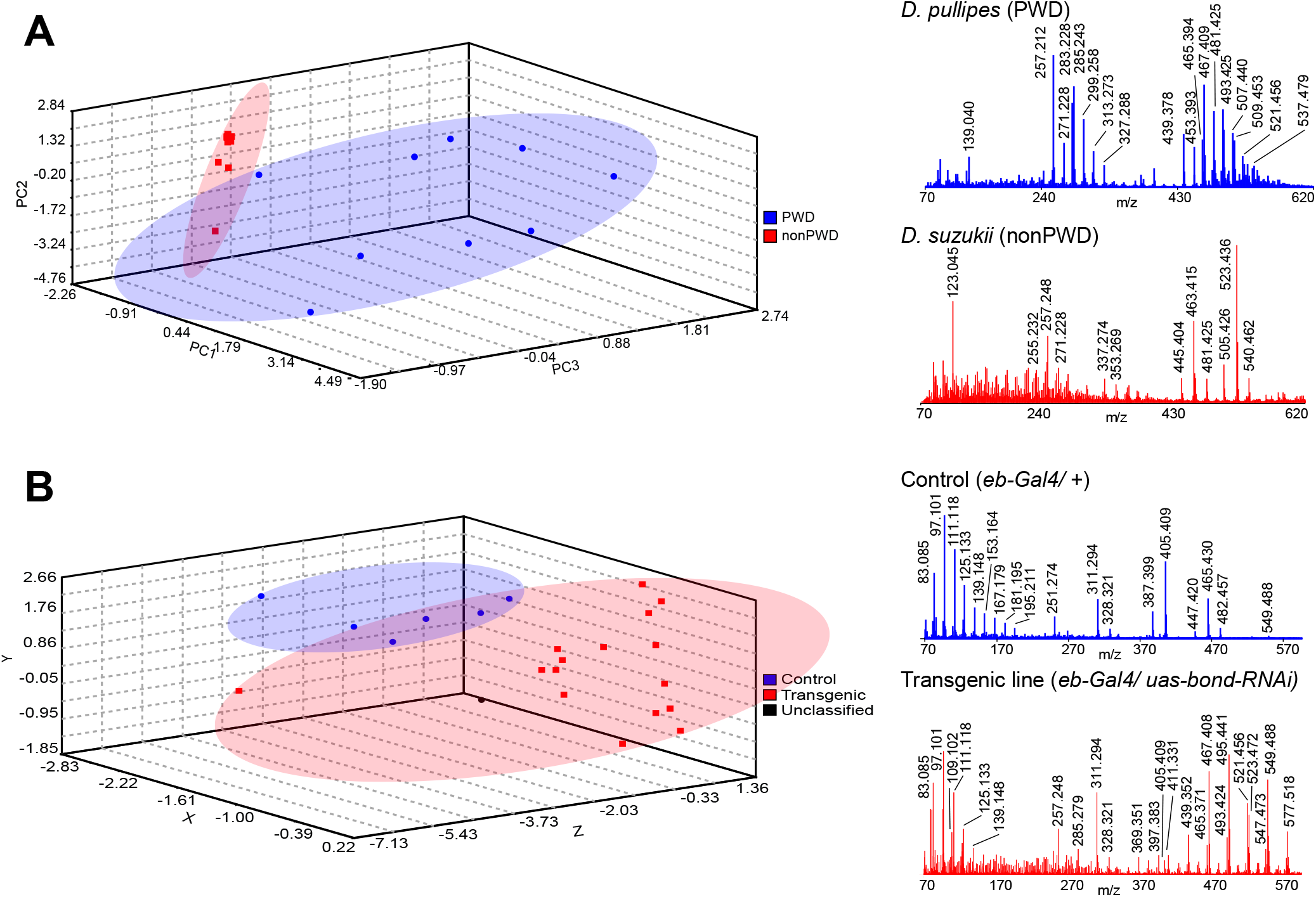
Classification by DART MS EB profiles. **(A)** Principal components analysis of the EB from 18 different drosophilid species differentiated between members of the Hawaiian Drosophila picture-wing subgroup (PWD) and nonPWD flies (LOOCV: 100%). Representative DART MS spectra reveal differences in chemical profile complexity from *D. pullipes* (PWD) and *D. suzukii* (nonPWD). **(B)** Linear discriminant analysis separated EB profiles of genetic controls from that of transgenic lines in which the expression of fatty acid-related genes in the EB have been genetically repressed (LOOCV: 96%). Representative DART MS spectra from a control genetic line (*bond-GAL4*/ +) and transgenic line (*bond-GAL4/ uas-bond-RNAi)* show the absence of CH503 at [M+H]^+^ 465.43 following repression of the gene *bond*, a very long chain fatty acid elongase ^20^. All EBs were measured at 300 °C.

We used the same clustering method to identify *D. melanogaster* transgenic lines with aberrant EB profiles. The ability to generate mutants and screen rapidly for phenotypic differences is a significant advantage of the *Drosophila* model system. We profiled the EBs of transgenic lines in which the expression of individual FA biosynthesis genes was genetically knocked down. Individuals with experimentally-induced changes in the EB lipid profile could be differentially separated from genetic controls using linear discriminant analysis with 92.5% accuracy (LOOCV; **Fig 4B**). One sample could not be classified. The results demonstrate that DART MS profiling of tissues is an effective method for the rapid screening of anomalous chemical profiles. This feature will allow mutants with defects in lipid metabolism to be swiftly identified.

## Conclusions

The direct analysis of tissue by DART MS generates rapid metabolite and lipid profiling from single pieces of tissue. This application will be particularly useful for samples that are rare or available in limited quantities. However, structural elucidation and distinguishing endogenous products from thermal decomposition artifacts will require a secondary form of MS confirmation. Both apolar and heavier, more polar molecules can be detected from single tissues with high reproducibility. In principal, quantitation with direct DART MS analysis of tissue is possible with the addition of an external standard although prior knowledge of the analyte concentration as well as its chemical properties will be required to select an appropriate standard with similar ionization properties. In addition, environmental contaminants, such as fatty acids, and signals resulting from in-source decomposition are commonplace and can be difficult to distinguish from endogenous products.

The ability to generate reproducible small molecule profiles based on relative abundances allows divergent phylogenetic lineages to be separated and phenotypic changes arising from genetic manipulation to be readily identified. The application of DART MS to tissue profiling will help to expand the “secretome” of molecules produced by pheromone glands as well as provide preliminary identification of natural products of interest produced within specialized compartments. Finally, the pairing of genetic screening with DART MS profiling can be extended towards metabolite and lipid profiling from single organs or biological compartments of other genetically tractable model organisms and applied to pharmaceutical studies for tissue-specific screening and detection of drug metabolites.

## Experimental

### Materials

All chemicals, unless otherwise stated, were purchased from Sigma-Aldrich (St. Louis, MO, USA).

### Animals

Hawaiian subgroup species *D. grimshawi, D. hemipeza* and, *D. silvestris* were obtained from Hawaiian Drosophila Research Stock Center and Insectary for Scientific Training and Advances in Research facility at the Univ. of Hawai‘i (https://icemhh.pbrc.hawaii.edu/instar/). Other picture-wing species (*D. pullipes, D. crucigera, D. adiastola, D. fasiculisetae, D. sproati*, and *D. ochracea* were collected from the Koke‘e State Park on Kaua‘i, the Nature Conservancy Waikamoi Preserve on Maui, South Kona Forest Reserve on Hawai‘i Island, and the Honouliuli Forest Reserve and O‘ahu. All other species were obtained from the Bloomington Drosophila Stock Center (Indiana, USA), National Drosophila Species Stock Center (Ithaca, NY, USA), and the Evolutionary Genetics Laboratory in Tokyo Metropolitan University (Japan). When possible, virgin males were used. Male reproductive status could not be confirmed with wild caught flies. Hawaiian flies were maintained on Clayton-Wheeler diet ^39^ at 19 °C and other species were raised at 25 °C on a 12 hours light/dark cycle on a standard diet of cornmeal, agar, dextrose, glucose, and 1.5% inactive yeast. Transgenic lines were generated using standard *Drosophila* genetic methods by crossing *bond-GAL4* females ^20^ to one the following *UAS-RNAi* lines from the Vienna Drosophila Resource Center: VDRC ID 46513, 23134, 1166, 7492, or 48662.

### Sample preparation for mass spectrometry

Single glands from males were dissected in phosphate buffered saline (PBS; 137 mM NaCl, 2.7 mM KCl, 10 mM Na2HPO4, 1.8 mM KH2PO4, pH 7.4) and placed directly on a stainless steel grid (DART QuickStrips; IonSense (IonSense LLC, Saugus, MA). For solvent studies, 1 µL of solvent was added to the tissue surface or an empty grid (for negative controls) immediately after dissection and allowed to evaporate. The following HPLC-grade solvents were tested: hexane, chloroform/ methanol (2:1 v/v), methanol, and ethanol. Five to seven replicates were prepared for each sample type. Immediately after dissection and placement on the sample grid, the strip was placed on a DART SVP linear rail system and moved through the ion source at 1 mm/sec at a distance of 0.5 cm from the inlet. Synthetic tributyrin, diolein (Cayman Chemical; Ann Arbor, MI, USA), *cis*-vaccenyl acetate (Cayman Chemical) and CH503 (previously synthesized ^40^) were prepared at 1 µM in hexane and 1 µL added to the sample grid.

### Direct tissue quantitation of tricosene

Five concentrations of tricosene diluted in hexane were tested: 0.0001, 0.001, 0.01, 0.1, and 1 mM. One µL of the standard was added to the EB immediately following placement on the grid. Five replicate EBs from 3 day old male flies were prepared for each concentration.

### Sample preparation for light microscopy

The EB from 3-4 day old males were dissected in PBS, fixed in 4 % paraformaldehyde for 10 min, then mounted on a glass slide in 50% glycerol. Digital Images were obtained with an Olympus BX51 microscope equipped with a Leica DFC 7000T color digital camera.

### Mass spectrometry and data analysis

Mass spectra were acquired with an atmospheric pressure ionization time-of-flight mass spectrometer (AccuTOF-DART 4G, JEOL USA, Inc., Peabody, MA) equipped with a DART SVP ion source (IonSense LLC, Saugus, MA) interface, placed 1 cm away from the sampling orifice. The instrument has a resolving power of 10,000 (FWHM definition) at *m/z* 500. Voltage settings and acquisition parameters for positive ion mode are as previously described ^35^. Briefly, the RF ion guide voltage was set at 500 V and the detector voltage set at 2200 V. The atmospheric pressure ionization interface potentials were as follows: orifice 1 = -15 or -21 V, orifice 2 = -5 V, ring lens = -5 V. Mass spectra were stored at a rate of one spectrum per second with an acquired *m/z* range of 100 – 1000. Calibration for exact mass measurements was accomplished by acquiring a mass spectrum of polyethylene glycol (average molecular weight 600) as an external reference standard in every data file.

### Fly-assisted laser desorption ionization mass spectrometry

Individual flies were attached to a custom-milled sample plate using double-sided tissue tape (7111, Louis Adhesive Tapes, Thailand). No matrix was used. Mass spectra were generated using a QSTAR Elite (AB Sciex, Toronto, CA) mass spectrometer equipped with a modified oMALDI2 ion source (AB Sciex) and a N_2_ laser (*λ*=337 nm) operated at a repetition rate of 40 Hz^9, 20^. Two mbar of N_2_ gas was used create the buffer gas environment for generation of ions. [M+K]^+^ potassium-bearing compounds constitute the major ion species. Spectra were internally calibrated to chitin signals at [M+K]^+^ 242.04, 648.20 and 851.28. Mass accuracy was ∼20 p.p.m. All spectra were acquired in positive ion mode and processed using MS Analyst software (Analyst QS 2.0, AB Sciex).

### Data analysis

The JEOL MassCenter software (v. 1.3.0.1) was used to calculate the area under the total or extracted ion chromatograms and to generate centroided profiles. Centroided profiles of each technical replicate were generated by summing signals within a 1 sec window of measurement. Preliminary determination of chemical categories is based on elemental composition prediction, predicted double bond equivalents, and isotope matching as performed with Mass Mountaineer (v. 5.1.13.0, RBC Software). For temperature studies, raw intensity counts for all centroided signals corresponding to a single chemical category were summed and normalized to the total area of all signals. For fragmentation studies, fragment percentage was calculated by dividing the signal abundance of the fragment (deacetylated molecule for pheromones; di-or monoglyceride for TAG and DAG standards) by the sum of the fragment and all adducts of the parent ion. For tricosene quantitation, the area beneath the extracted ion chromatogram for tricosene ([M+H]^+^ 323.36) was used to calculate abundance. For solvent studies, the area of the extracted ion chromatograms corresponding to each chemical category was normalized to the area beneath the total ion chromatogram. The relative abundances from 5-7 replicates were summed and averaged. Background levels of FAs were established by averaging the absolute abundances from 5 replicate measurements. Statistical analysis was performed with Prism v.9.2 (GraphPad Software, San Diego, CA).

### Classification analysis

Centroided mass spectra were exported as text files using MassCenter for classification and analysis by Mass Mountaineer. For linear discriminant analysis (LDA) and kernel principal component analysis (KPCA), a signal threshold of 10% was used to extract features from centroided mass spectral data. Five *m/z* values (for LDA) and 11 *m/z* values (for KPCA) with a mass tolerance of 10 mmu were used in the training set. The normalized relative abundance of each of the features was calculated relative to the highest intensity peak for the spectrum. Five to seven biological replicates for each species or genotype were used for classification. Predictive models were tested with leave on out cross validation (LOOCV).

## Figure legends

**Supp. Table 1.** Signals and chemical classes identified from the direct analysis of the EB using DART MS and laser desorption ionization MS.

**Supp. Fig. 1.** (**A-F**) DART MS spectra measured from six replicate EBs analyzed on the same sample strip at 300 °C. The spectra are qualitatively similar and contain the major signals corresponding to cVA, deacetylated cVA, CH503, and deacetylated CH503. Some quantitative variation is evident - e.g., in **B** and **F**, the intensity of the FA signal at m/z 257.248 is comparable or higher than that of cVA and CH503. Unless otherwise indicated, ions are detected as [M+H]^+^.

**Supp. Fig. 2.** DART MS analysis of cVA and CH503 synthetic standards shows a loss of acetate from both molecules and loss of water from CH503. The patterns of pheromone decomposition are consistent with those observed from direct analysis of EB tissue. Signals found in both synthetic standards and direct tissue analysis are labeled in blue. Analysis was performed at 300 °C.

**Supp. Fig. 3.** Effect of ion source temperature on synthetic tributyrin [(4:0/ 4:0/ 4:0)-TAG] and diolein [(18:1/ 18:1)-DAG]. (**A**) The loss of C4:0 acyl groups from tributyrin significantly increases with temperature; *p=0.03-0.04; **: p=0.0051; ***: p=0.0002-0.0007; ****: p<0.0001, Kruskal-Wallis. **(B)**. DAGs are detected as dehydrated ions at every temperature tested; **: p=0.0023, Kruskal-Wallis. For both graphs, each point represents the mean ± S.D, n=5-7.

**Supp. Fig. 4.** Representative spectra from EBs washed with various solvents; NS: no solvent. Mass signals corresponding to cVA, CH503, deacetylated fragments, fatty acids (FA), diacylglycerides (DAGs) and triacylglycerides (TAGs) are labeled.

**Supp. Fig. 5.** The abundance of fatty acids (FA) in solvent-only controls and EBs treated with solvents measured at 250 °C. The FA levels were similar between solvent-only and solvent-treated EBs in ethanol and chloroform/ MeOH (CM) conditions. Levels of FAs in untreated EBs or EBs treated with MeOH or hexane could be distinguished from baseline levels. For all graphs, bars indicate mean ± S.E.M; Mann-Whitney test, n=4-6.

## Author Contributions

NC: methodology and investigation.; JYY: Conceptualization, methodology, investigation, validation, formal analysis, resources, and writing – original draft, review, and editing.

There are no conflicts to declare.

## Acknowledgements

The DART MS instrument was acquired through a grant from the Department of Defense, U.S. Army Research Office (W911NF1610216 to J.Y.Y). The work is supported by the National Institutes of Health (Grant No. P20GM125508), Hawai‘i Community Foundation (19CON-95452), and the National Science Foundation (Grant No. 20256669). This article reports data obtained at the University of Hawaiʻi at Mänoa Microbial Genomics and Analytical Laboratory Core which is supported by the National Institutes of Health (P20GM125508). We thank Kenneth Kaneshiro, Kelvin Kanegawa, Matthew J. Medeiros, Donald Price, Kelli Konicek and the Hawaiian Drosophila Research Stock Center for providing fly samples.

## Notes

### Competing Interest Statement

The authors have declared no competing interest.

