## Supplemental Data for "In-situ lipid profiling of insect pheromone glands by Direct Analysis in Real Time Mass Spectrometry"

**Supp. Table 1.** Signals and chemical classes identified from the direct analysis of the EB using DART MS and laser desorption ionization MS.

| Elemental composition[1] | Meas. *m/z* | Calc. *m/z* | Δ (mmu)[2] | Ion Form | #C: #DB[3] | Chemical class[4] | LDI MS[5] *m/z* | Calc. *m/z* | Δ (mmu) |
|---|---|---|---|---|---|---|---|---|---|
| C14 H25 O2 | 225.182 | 225.185 | -0.003 | $[M+H]^+$ | C14:2 | FA | - | - | - |
| C14 H27 O2 | 227.203 | 227.201 | 0.002 | $[M+H]^+$ | C14:1 | FA | 265.16 | 265.16 | 0.00 |
| C14 H29 O2 | 229.213 | 229.217 | -0.004 | $[M+H]^+$ | C14:0 | FA | 267.16 | 267.17 | -0.01 |
| C18 H34 | 251.274 | 251.274 | 0.000 | $[M+H]^+$ | - | cVA-OAc | - | - | - |
| C16 H31 O2 | 255.232 | 255.232 | 0.000 | $[M+H]^+$ | C16:1 | FA | 293.18 | 293.19 | -0.01 |
| C16 H33 O2 | 257.247 | 257.248 | -0.001 | $[M+H]^+$ | C16:0 | FA | 295.20 | 295.20 | - |
| C18 H33 O2 | 281.246 | 281.248 | -0.002 | $[M+H]^+$ | C18:2 | FA | 319.20 | 319.20 | - |
| C18 H35 O2 | 283.259 | 283.264 | -0.005 | $[M+H]^+$ | C18:1 | FA | 321.21 | 321.22 | -0.01 |
| C26 H34 O1 N1 | 376.269 | 376.264 | 0.005 | $[M+H]^+$ | - | unk | - | - | - |
| C20 H39 O2 | 311.294 | 311.295 | -0.001 | $[M+H]^+$ | - | cVA | 349.25 | 349.25 | 0.00 |
| C22 H41 O4 | 369.299 | 369.300 | -0.001 | $[M+H]^+$ | - | unk | 407.25 | 407.26 | -0.01 |
| C28 H53 O1 | 405.409 | 405.409 | 0.000 | $[M+H]^+$ | - | CH503-OAc | 443.35 | 443.37 | -0.02 |
| C30 H57 O3 | 465.43 | 465.431 | -0.001 | $[M+H]^+$ | - | CH503 | 503.37 | 503.39 | -0.02 |
| C33 H61 O4 | 521.456 | 521.457 | -0.001 | $[M+H]^+$ | C30:1 | DAG-$H_2O$ | - | - | - |
| C33 H63 O4 | 523.471 | 523.470 | 0.001 | $[M+H]^+$ | C30:0 | DAG-$H_2O$ | 561.43 | 561.43 | 0.00 |
| C33 H63 O5 | 539.462 | 539.468 | -0.006 | $[M+H]^+$ | C30:1 | DAG | 577.42 | 577.42 | 0.00 |
| C33 H65 O5 | 541.488 | 541.483 | 0.005 | $[M+H]^+$ | C30:0 | DAG | - | - | - |
| C35 H61 O4 | 545.455 | 545.457 | -0.002 | $[M+H]^+$ | C32:3 | DAG-$H_2O$ | - | - | - |
| C35 H63 O4 | 547.472 | 547.469 | 0.003 | $[M+H]^+$ | C32:2 | DAG-$H_2O$ | - | - | - |
| C35 H63 O5 | 563.47 | 563.468 | 0.002 | $[M+H]^+$ | C32:3 | DAG | - | - | - |
| C35 H65 O4 | 549.488 | 549.488 | 0.000 | $[M+H]^+$ | C32:1 | DAG-$H_2O$ | 587.44 | 587.44 | 0.00 |
| C35 H65 O5 | 565.483 | 565.483 | 0.000 | $[M+H]^+$ | C32:2 | DAG | 603.43 | 603.44 | -0.01 |
| C35 H65 O6 | 581.475 | 581.478 | -0.003 | $[M+H]^+$ | C32:1 | TAG | - | - | - |
| C35 H67 O5 | 567.498 | 567.499 | -0.001 | $[M+H]^+$ | C32:0 | DAG | 605.44 | 605.45 | -0.01 |
| C37 H65 O4 | 573.492 | 573.488 | 0.004 | $[M+H]^+$ | C34:3 | DAG-$H_2O$ | - | - | - |
| C37 H67 O4 | 575.501 | 575.504 | -0.003 | $[M+H]^+$ | C34:2 | DAG-$H_2O$ | - | - | - |
| C37 H67 O5 | 591.5 | 591.499 | 0.001 | $[M+H]^+$ | C34:3 | DAG | - | - | - |
| C37 H69 O4 | 577.519 | 577.52 | -0.001 | $[M+H]^+$ | C34:1 | DAG-$H_2O$ | 615.47 | 615.48 | -0.01 |
| C37 H69 O5 | 593.514 | 593.515 | -0.001 | $[M+H]^+$ | C34:2 | DAG | - | - | - |
| *C37 H72 O5 N1* | *610.536* | *610.541* | *-0.005* | $[M+NH4]^+$ | *C34:2* | *DAG* | - | - | |
| C37 H71 O5 | 595.522 | 595.53 | -0.008 | $[M+H]^+$ | C34:1 | DAG | 633.47 | 633.48 | -0.01 |
| C39 H69 O4 | 601.52 | 601.52 | 0.000 | $[M+H]^+$ | C36:3 | DAG-$H_2O$ | - | - | - |
| C39 H71 O4 | 603.535 | 603.53 | 0.005 | $[M+H]^+$ | C36:2 | DAG-$H_2O$ | - | - | - |
| C39 H71 O5 | 619.53 | 619.53 | 0.000 | $[M+H]^+$ | C36:3 | DAG | - | - | - |
| C39 H73 O4 | 605.551 | 605.551 | 0.000 | $[M+H]^+$ | C36:1 | DAG-$H_2O$ | - | - | - |
| C40 H77 O4 | 621.576 | 621.582 | -0.006 | $[M+H]^+$ | C37:0 | DAG-$H_2O$ | 659.52 | 659.53 | -0.01 |
| C41 H82 O6 N1 | *684.606* | 684.614 | -0.008 | $[M+NH4]^+$ | C38:0 | TAG | - | - | - |
| C43 H86 O6 N1 | *712.641* | 712.646 | -0.005 | $[M+NH4]^+$ | C40:0 | TAG | - | - | - |
| C43 H84 O6 N1 | *710.612* | 710.609 | 0.003 | $[M+NH4]^+$ | C40:1 | TAG | - | - | - |
| C45 H87 O6 | 723.646 | 723.65 | -0.004 | $[M+H]^+$ | C42:0 | TAG | 761.59 | 761.60 | -0.01 |
| *C45 H90 O6 N1* | *740.673* | *740.677* | *-0.004* | $[M+NH4]^+$ | *C42:0* | *TAG* | - | - | - |
| C45 H85 O6 | 721.631 | 721.635 | -0.004 | $[M+H]^+$ | C42:1 | TAG | 759.58 | 759.59 | -0.01 |
| *C45 H88 O6 N1* | *738.654* | *738.661* | *-0.007* | $[M+NH4]^+$ | *C42:1* | *TAG* | - | - | - |
| C47 H94 O6 N1 | 768.7 | 768.708 | -0.008 | $[M+NH4]^+$ | C44:0 | TAG | - | - | - |
| C47 H89 O6 | 749.663 | 749.666 | -0.003 | $[M+H]^+$ | C44:1 | TAG | 787.60 | 787.62 | -0.02 |
| *C47 H92 O6 N1* | *766.687* | *766.692* | *-0.005* | $[M+NH4]^+$ | *C44:1* | *TAG* | - | - | - |
| C47 H87 O6 | 747.645 | 747.65 | -0.005 | $[M+H]^+$ | C44:2 | TAG | 785.58 | 785.60 | -0.02 |
| *C47 H90 O6 N1* | *764.665* | *764.677* | *-0.012* | $[M+NH4]^+$ | *C44:2* | *TAG* | - | - | - |
| C49 H93 O6 | 777.694 | 777.697 | -0.003 | $[M+H]^+$ | C46:1 | TAG | 815.64 | 815.65 | -0.01 |
| *C49 H96 O6 N1* | *794.718* | *794.723* | *-0.005* | $[M+NH4]^+$ | *C46:1* | *TAG* | - | - | - |
| C49 H91 O6 | 775.681 | 775.682 | -0.001 | $[M+H]^+$ | C46:2 | TAG | 813.62 | 813.64 | -0.02 |
| *C49 H94 O6 N1* | *792.702* | *792.708* | *-0.006* | $[M+NH4]^+$ | *C46:2* | *TAG* | - | - | - |
| C49 H89 O6 | 773.661 | 773.666 | -0.005 | $[M+H]^+$ | C46:3 | TAG | - | - | - |
| *C49 H92 O6 N1* | *790.688* | *790.692* | *-0.004* | $[M+NH4]^+$ | *C46:3* | *TAG* | - | - | - |
| C51 H97 O6 | 805.724 | 805.729 | -0.005 | $[M+H]^+$ | C48:1 | TAG | 843.66 | 843.68 | -0.02 |
| *C51 H100 O6 N1* | *822.748* | *822.755* | *-0.007* | $[M+NH4]^+$ | *C48:1* | *TAG* | - | - | - |
| C51 H95 O6 | 803.71 | 803.713 | -0.003 | $[M+H]^+$ | C48:2 | TAG | 841.68 | 841.67 | 0.01 |
| *C51 H98 O6 N1* | *820.731* | *820.739* | *-0.008* | $[M+NH4]^+$ | *C48:2* | *TAG* | - | - | - |
| C51 H93 O6 | 801.694 | 801.698 | -0.004 | $[M+H]^+$ | C48:3 | TAG | - | - | - |

[1]Italics indicate molecules that were detected as $[M+H]^+$ and $[M+NH_4]^+$; [2]mmu: milli-mass units; [3]No. of carbons and no. of double bonds predicted in FA chains; [4]Assignment is based on exact mass measurements and degree of unsaturation; some FAs and DAGs may be decomposition products; FA: fatty acid; DAG: diacylglyceride; TAG: triacylglyceride; unk: unknown; [5]LDI: laser desorption ionization MS; signals are detected as $[M+K]^+$.

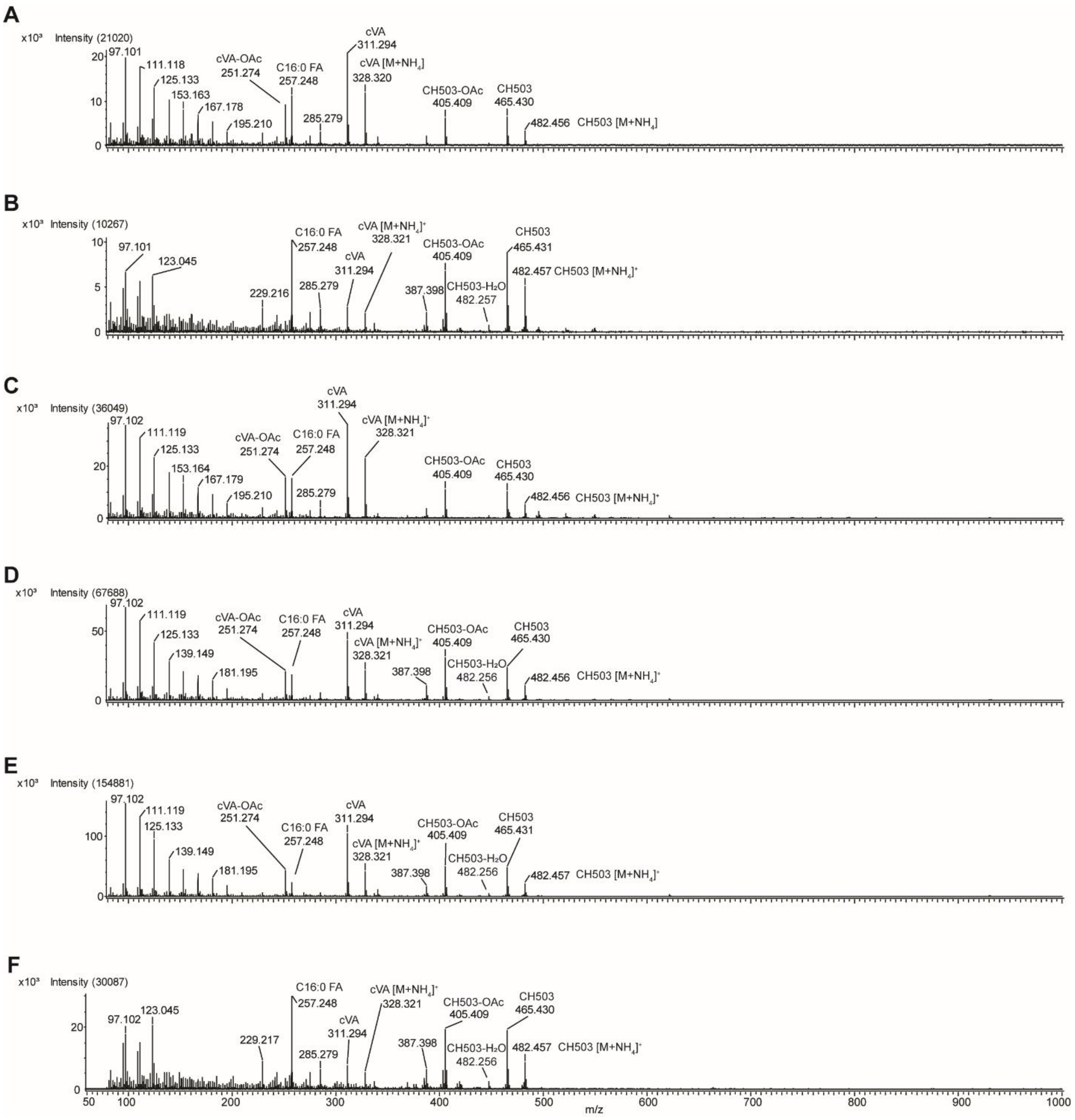


**Supp. Fig. 1.** (**A-F**) DART MS spectra measured from six replicate EBs analyzed on the same sample strip at 300 ˚C. The spectra are qualitatively similar and contain the major signals corresponding to cVA, deacetylated cVA, CH503, and deacetylated CH503. Some quantitative variation is evident - e.g., in **B** and **F**, the intensity of the FA signal at m/z 257.248 is comparable or higher than that of cVA and CH503. Unless otherwise indicated, ions are detected as $[M+H]^+$.

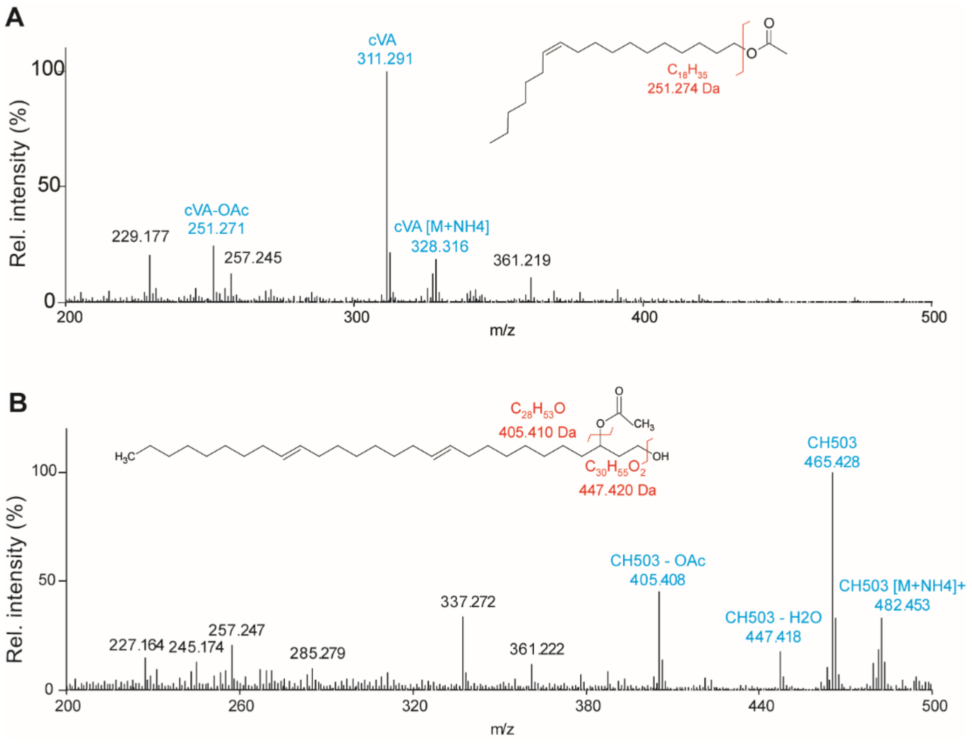


**Supp. Fig. 2.** DART MS analysis of cVA and CH503 synthetic standards shows a loss of acetate from both molecules and loss of water from CH503. The patterns of pheromone decomposition are consistent with those observed from direct analysis of EB tissue. Signals found in both synthetic standards and direct tissue analysis are labeled in blue. Analysis was performed at 300 ˚C.

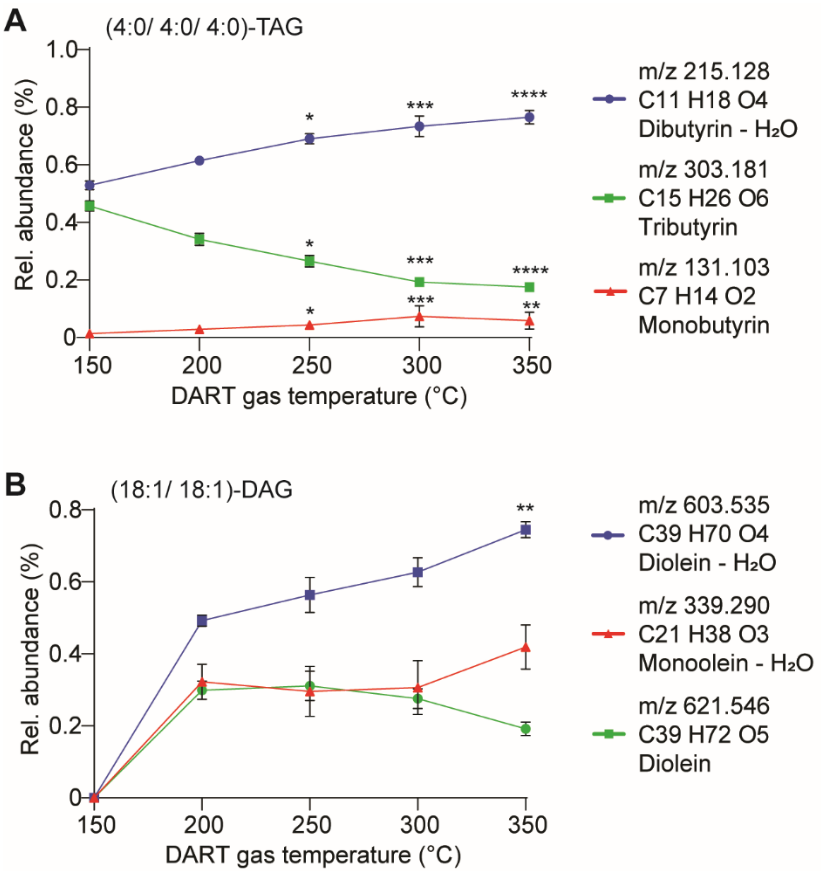


**Supp. Fig. 3.** Effect of ion source temperature on synthetic tributyrin [(4:0/ 4:0/ 4:0)-TAG] and diolein [(18:1/ 18:1)-DAG]. (**A**) The loss of C4:0 acyl groups from tributyrin significantly increases with temperature; *p=0.03-0.04; **: p=0.0051; ***: p=0.0002-0.0007; ****: p<0.0001, Kruskal-Wallis. **(B)**. DAGs are detected as dehydrated ions at every temperature tested; **: p=0.0023, Kruskal-Wallis. For both graphs, each point represents the mean ± S.D, n=5-7.

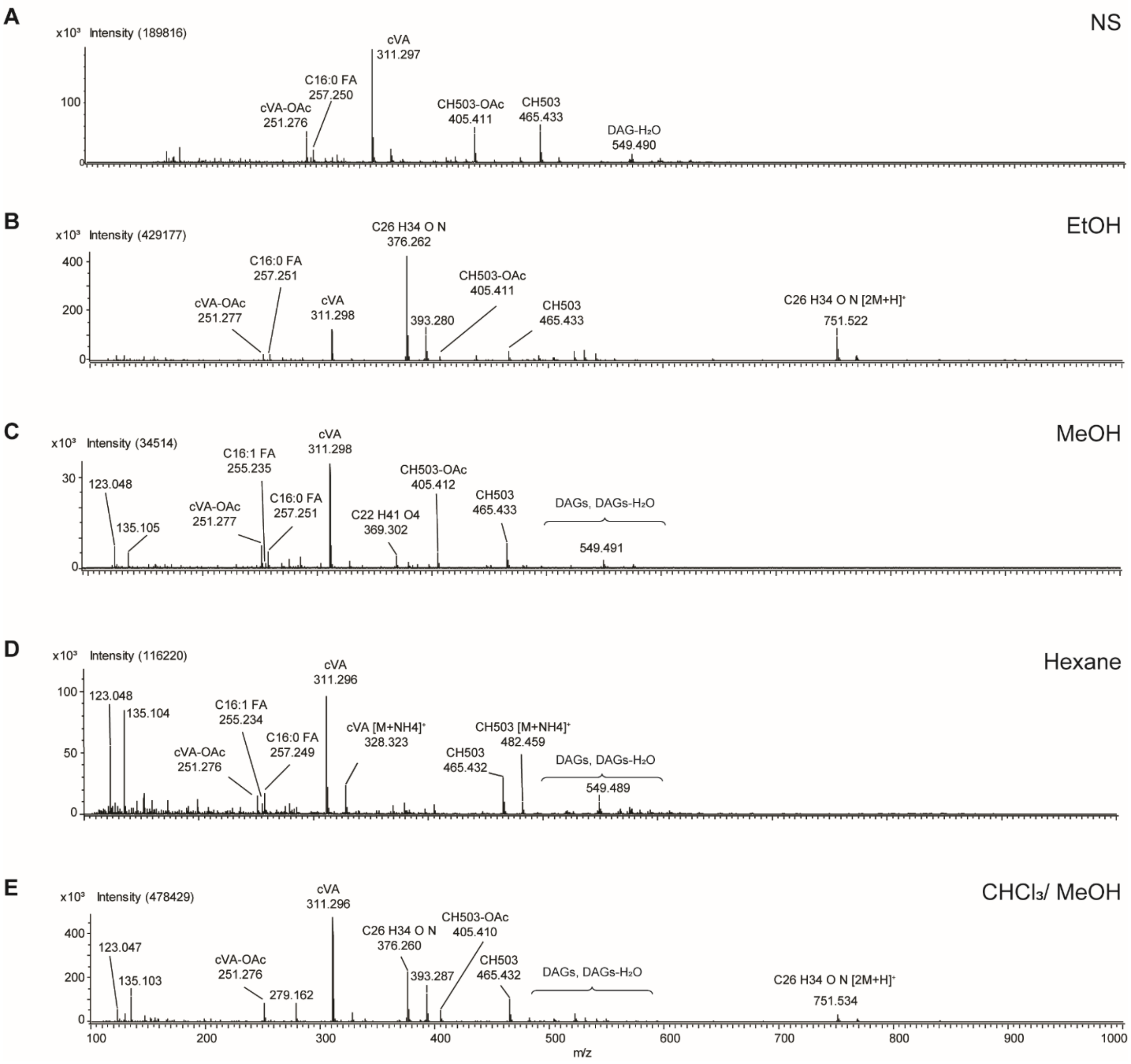


**Supp. Fig. 4.** Representative spectra from EBs washed with various solvents; NS: no solvent. Mass signals corresponding to cVA, CH503, deacetylated fragments, fatty acids (FA), diacylglycerides (DAGs) and triacylglycerides (TAGs) are labeled.

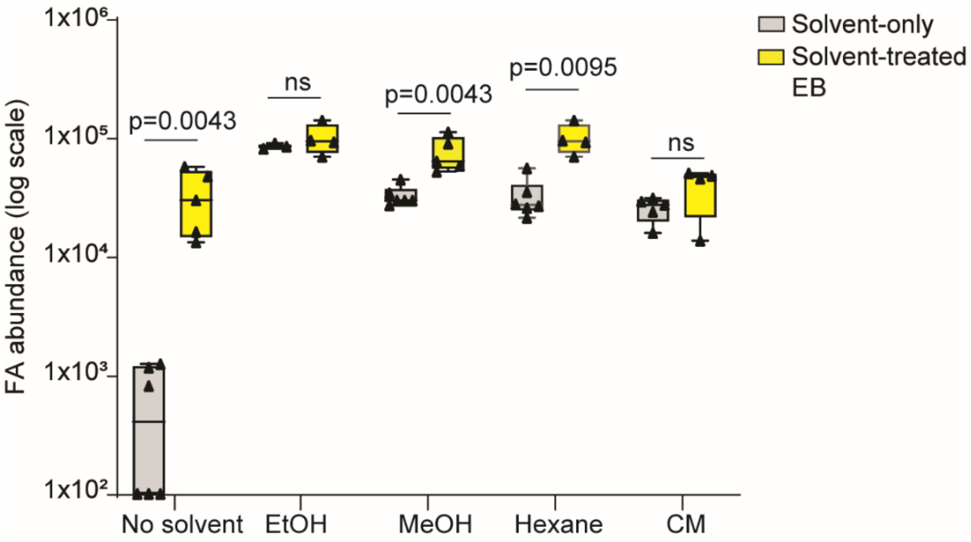


**Supp. Fig. 5.** The abundance of fatty acids (FA) in solvent-only controls and EBs treated with solvents measured at 250 ˚C. The FA levels were similar between solvent-only and solvent-treated EBs in ethanol and chloroform/ MeOH (CM) conditions. Levels of FAs in untreated EBs or EBs treated with MeOH or hexane could be distinguished from baseline levels. For all graphs, bars indicate mean ± S.E.M; Mann-Whitney test, n=4-6.
